# Mass Spectrometric Characterization of Narcolepsy-Associated Pandemic 2009 Influenza Vaccines

**DOI:** 10.1101/2020.08.21.256180

**Authors:** Aditya Ambati, Guo Luo, Elora Pradhan, Jacob Louis, Ling Lin, Ryan Lieb, Hanna Maria Ollila, Thomas Poiret, Christopher Adams, Emmanuel Mignot

**Affiliations:** Stanford Center for Sleep Sciences and Medicine, Department of Psychiatry and Behavioral Sciences, Stanford University, 3165 Porter Drive, Stanford CA, USA; Stanford Mass Spectrometry Core, 333 Campus Drive, Mudd 175, Stanford University, Stanford, CA, USA; Department of Laboratory Medicine, Karolinska Institutet, Stockholm, Sweden

**Keywords:** narcolepsy, influenza vaccine, Mass Spectrometry, mutations

## Abstract

The onset of narcolepsy, an irreversible sleep disorder, has been associated with 2009 influenza pandemic (pH1N1) infections in China, and with ASO3-adjuvanted pH1N1 vaccinations using Pandemrix in Europe. Intriguingly, however, the increased incidence was only observed following vaccination with Pandemrix but not Arepanrix in Canada. In this study, the mutational burden of actual vaccine lots of Pandemrix (n=6) and Arepanrix (n=5) sourced from Canada, and Northern Europe were characterized by mass spectrometry. The four most abundant influenza proteins across both vaccines were nucleoprotein NP, hemagglutinin HA, matrix protein M1, with the exception that Pandemrix harbored a significantly increased proportion of neuraminidase NA (7.5%) as compared to Arepanrix (2.6%). Most significantly, 17 motifs in HA, NP, and M1 harbored mutations, which significantly differed in Pandemrix versus Arepanrix. Among these, a 6-fold higher deamidation of HA146 (N to D) in Arepanrix was found relative to Pandemrix, while NP257 (T to A) and NP424 (T to I) were increased in Pandemrix. DQ0602 binding and tetramer analysis with mutated epitopes were conducted in Pandemrix-vaccinated cases versus controls but were unremarkable. Pandemrix harbored lower mutational burden than Arepanrix, indicating higher similarity to wild-type 2009 pH1N1, which could explain differences in narcolepsy susceptibility amongst the vaccines.

## Introduction

Type 1 Narcolepsy (T1N) is a disabling disorder characterized by excessive daytime sleepiness, irresistible daytime sleep attacks, and sudden episodes of loss of muscle tone following emotions such as laughter, a symptom known as cataplexy^1^. Genetic and immunological studies have shown that the disorder is autoimmune, and likely mediated by T cell attacks targeting hypocretin neurons, a population of 20,000 neurons located in the posterior hypothalamus^2,3^. Hypocretin peptides (hypocretin-1 and 2), also called orexins, are homologous peptides derived from a single precursor, pre-pro-hypocretin. The precursor undergoes cleavage, with cleaved products undergoing post-translational modification, notably C-terminal amidation, a transformation indispensable to biological activity. Hypocretins are critical regulators of wakefulness and Rapid Eye Movement sleep (REM sleep), and lack of hypocretin transmission is causal to the symptoms of the disorder^4^.

Until recently, the rationale for an autoimmune basis for narcolepsy was based mainly on epidemiological and genetic evidence. First, there is a uniquely strong association between narcolepsy and 6p21.3, a region of the genome, including the HLA locus^5–7^. More specifically, 97% of narcoleptic patients carry at least one copy of HLA DQB1*06:02 (DQ0602) across ethnicities, an HLA class II allele found in 25% of the general population^5^. Additional HLA effects include trans modulatory effects of other DQ molecules, weak effects in HLA class I, notably a trans-ethnic association with HLA-A*11:01^8^ and an impact of DQB1*03:01 on the age of onset^9^. Besides, genome-wide association studies have found that narcolepsy is associated with T-cell receptor loci TRA & TRB, and immune genes such as CTSH, P2RY11, ZNF265, IFNAR1, and TNSF4^10–13^. As all these loci are involved in immune regulation and other autoimmune diseases, an autoimmune mediation of hypocretin cell death has long been proposed as the cause of narcolepsy. Of notable interest is the fact the TCR loci associated with narcolepsy are modulators of TRAJ24 and TRBV4-2, TCR segments only involved in 0.8% and 0.7% of the total TCR repertoire, respectively. Another notable finding is an association with an INFAR1 polymorphism that maps to a quantitative trait loci (QTL) for dendritic cell reactivity to flu infection^13^.

Narcolepsy studies have described environmental triggers in addition to the genetic susceptibilities; specifically, studies have noted increased humoral (IgG) and cellular (IFNγ) responses to streptococcus pyogenes infection^14,15^. Similarly, epidemiological studies have suggested increased frequency of strep infections and flu-like illness in patients before developing narcolepsy^16^. Most recently, the data has most strongly implicated influenza-A infection and vaccination. Following the 2009-2010 H1N1 “swine flu” influenza pandemic infection in China, increased T1N onsets were observed^11^. In European countries, a significant 4-16-fold increase in the risk of developing narcolepsy in children was observed a few months following an aggressive pandemic H1N1 (pH1N1) flu vaccination campaign with the vaccine Pandemrix^17^. This phenomenon was first reported in Finland and Sweden, where coverage with Pandemrix was respectively 50% and 60%, but later confirmed in almost all countries where it was used (Ireland, England, France, Norway)^18–24^. In these cases, Pandemrix increased the incidence of narcolepsy by a factor of 2-15, from around 1/150,000 to 1/15,000 cases per year in children. The risk was mostly increased in younger children, but there was still a significant, albeit weaker effect in adults^20^. Similar increases in narcolepsy incidence were not observed in countries where other pandemic vaccines, notably Arepanrix in Canada, were used, thereby elucidating the impression that Pandemrix uniquely triggered narcolepsy^25^.

Starting in May 2009, when pandemic H1N1 first appeared in Mexico, vaccine manufacturers began to plan the production of a specialized vaccine targeting the new strain for vaccination the following winter, a concise timeline. The creation of vaccine strains involves growing strains derived from pathogenic strains reassorted with PR8 (an old 1918-H1N1-like strain 08/35 from Puerto Rico) in eggs. The reassortant strain is typically constructed by the New York Medical Center (NYMC), which is then distributed to manufacturers for growing millions of doses in eggs in specialized factories. The NYMC H1N1-like vaccine strains produced for the 2009-2010 swine flu campaign used A/H1N1/California/7/2009 as the pathogenic strain, so that only Hemagglutinin (HA), Neuraminidase (NA), and polymerase PB1 are derived from A/H1N1/California/7/2009, while other proteins are PR8 derived. In close succession, NYMC-X-179A and NYMC X-181, a higher growth reassortant derived from X-179A were created, with the former strain having been used more widely (X-181 was only used toward the end of the season in some cases)^26^. Once distributed, vaccine manufacturers used their own patented process to produce vaccines using X-179A and X-181. In egg-based vaccine production processes, candidate vaccine viruses are grown in eggs per current FDA regulatory requirements. To do so, X-179A is injected into fertilized hen’s eggs and incubated for several days to allow viruses to replicate. The virus-containing allantoic fluid is then harvested from the eggs, viruses inactivated (killed), and virus antigens purified, with the general goal of preferentially isolating viral surface proteins HA and NA, which are most important for protective antibody responses sought with vaccine administration^27,28^.

Using this process, the manufacturer GlaxoSmithKline created Pandemrix and Arepanrix, both AS03-adjuvanted vaccines; the AS03 adjuvant is an immunological agent added to the vaccine to boost the immune system’s response to the target antigen while reducing the dosage of the viral antigen (antigen sparing, a property that was desirable considering short time of production)^28^. Arepanrix was produced and used in Canada around the same time that Pandemrix was deployed in Europe, but it did not sharply increase narcolepsy risk in Quebec^25,29^. This is notable as Arepanrix is almost identical to Pandemrix with a same adjuvant from the same geographic origin as well as similar viral composition. However, different viral antigen purification techniques were used for either vaccine -- Fluarix for Pandemrix and Flulaval for Arepanrix^30^. Why Pandemrix in Europe and not Arepanrix in Canada triggered narcolepsy cases is unknown. One possibility may be the differential composition of the vaccine, notably due to the fact the viral antigens were extracted using different manufacturing processes. Another possibility involves the presence of other factors differentiating Canada and Europe at the time of vaccination. After all, it is essential to note that even with Pandemrix, only 1/16,000 vaccinated children (or 1/4000 DQB0602 positive subjects) developed narcolepsy^19^, so that almost undoubtedly other environmental or stochastic factors are involved in addition to vaccine trigger and genetic background. As Canadian and northern European populations are similar in term of DQ0602 frequency (and other narcolepsy-associated genetic factors), a possibility could be a differential immune history of both population regarding past flu or the fact that in Northern Europe vaccination occurred shortly before or exactly when the pandemic H1N1 infection affected the population. At the same time, in Canada the bulk of vaccination occurred immediately after the pandemic flu started to change the population^23^.

Our understanding of narcolepsy immunology changed significantly a few months ago, thanks to two studies ^31,32^. Latorre et al. used an ultrasensitive technique involving polyclonal expansion and cloning of CD45RA−CD4+ memory T cells as lines, followed by the screening of these lines for reactivity as a surrogate of proliferation to autoantigen peptide pools presented by autologous B cells. Screening Peripheral Blood monocyte cells (PBMCs) of 19 T1N cases (15 with documented hypocretin deficiency, defined by low hypocretin-1 in the cerebrospinal fluid) and 12 controls, these authors found strong line reactivity to HCRT in all patients versus no or limited responses in 12 DQ0602 controls, with significantly higher reactivity in T1N. These authors also screened these same cell lines for proliferative reactions to seasonal Influenza A antigens and found responses to be comparable in patients and controls, concluding that flu antigen mimicry could not be detected in these autoreactive cells, which were mostly DR rather than DQ restricted^31^. On other hand, Luo et al^32^ screened peptides derived from HCRT and flu strains including pH1N1 for DQ0602 binding and presence of cognate tetramer-peptide specific CD4+ T cells in 35 T1N cases and 22 DQ0602 controls finding higher reactivity to influenza pHA^273-287^ (pH1N1 specific) and C-amidated but not native version of HCRT^54-66^ and HCRT^86-97^ sequences (two homologous sequences we denoted HCRT^NH2^) in T1N when presented by DQ0602. Previous sequence homology between pHA^273-287^ and HCRT^54-66^ or HCRT^86-97^ had been noted before, suggesting possible molecular mimicry. Most interestingly, T cell receptors isolated from cells reacting to these peptides contained two specific TRAJ24 and TRBV 4-2 segments in increased abundance, suggesting molecular mimicry to pHA^273-287^ and T cell autoreactivity to HCRT^NH2^ is causal to narcolepsy. However, some reactivity was also found in DQ0602 controls. This second paper, with its links to the genetic predisposition to narcolepsy, strongly suggests that presentation by DQ0602 of HCRT^NH2^ is causal to narcolepsy pathophysiology. It also identifies pHA^273-287^ as a likely mimic of HCRT^NH2^, but of course, does not exclude the possibility that other sequence mimics are present in different strains of flu, as narcolepsy has been linked to seasonal variation before 2009-2010^11^. The recent findings offer the opportunity of identifying a whole array of cross-reacting sequences that may play a role in flu-triggering of narcolepsy.

Relatively few studies have examined composition differences across flu vaccines such as Pandemrix and Arepanrix. Comparative studies of antibody reactivity to Pandemrix and Arepanrix antigens in both post-Pandemrix patients and control individuals found that post-Pandemrix-vaccinated children had poorer antibody reactivity to Arepanrix, suggesting antigenic differences in antibody determinants^33^. This could be important, although one would expect that differential susceptibility would be more likely due to T cell response differences. High-resolution gel electrophoresis quantitation and Mass Spectrometry (MS) identification analyses revealed higher amounts of structurally altered viral nucleoprotein (NP) in Pandemrix versus Arepanrix, a finding that can also be noted in a 2-D gel study by Jacob et al^30^. These results suggest complex protein aggregate conformation differences that could be relevant to differential activity of these vaccines, most notably in terms of antibody response to NP. Jacob et al^30^ comparing single batches of Pandemrix, Arepanrix and Focetria (a Novartis subunit-specific vaccine that also used a squalene adjuvant and was enriched in HA) found that viral proteins NP and NA as well as selected non-viral chicken proteins (PDCD6IP, TSPAN8, H-FABP, HSP, and TUB) were more abundant in Pandemrix compared to Arepanrix. The study also found an accumulation of a specific mutation in Arepanrix, 146N to D in Arepanrix, but as the batch studied had only been synthesized in 2010 and had never been used for vaccination, it was hard to be sure if this mutation had been present in earlier batches. Finally, Ahmed et al^34^ found an increased quantity of cross-reactive antibodies to structurally altered NP epitope NP116I with an epitope from HCRTR2 protein in narcolepsy cases. Although the proportion of NP116I mutations in both Arepanrix and Pandemrix was similar and, therefore, unlikely to be a causal effect^35^, increased cross-reactive antibodies between NP116I and HCRTR2 were not observed in other studies^36^.

In the current study, we extended the previous findings from Jacob et al. .^30^ by characterizing the protein content and mutational burden in 6 Pandemrix lots and 5 Arepanrix lots in relation to X-179A, the influenza strain from which both vaccines were derived. Data shows batch diversity in mutation burden that could be relevant to vaccine responses in some cases but not a distinct difference that could explain why Pandemrix was more associated with narcolepsy onsets than Arepanrix. Mutational drift and low-level mutations are present in some vaccine batches (as occurring in wild type virus), and this should be taken into consideration when studying the effects of these vaccines in the population.

## Methods

### Vaccines

Pandemrix doses used in the 2009 vaccination campaign were sourced from Sweden (3), France (1), and additionally provided by GSK (2), while Arepanrix was sourced from Canada (4) and supplied by GSK (1). All except for two lots were monovalent bulks (see Table 1). The vaccines were all derived from the X-179A, a vaccine strain that built on the PR8 backbone and was populated with pH1N1 proteins (HA, NA and PB1) from the A/California/07/2009 strain. Pandemrix lots were prepared using the Fluarix process in Dresden while the Arepanrix lots were processed using the Flulaval process in Saint Foye as described^30^. All the vaccines were stored at +4C until assayed.

**Table 1:** Characteristics of Pandemrix and Arepanrix lots used in the MS characterization

| **Vaccine** | **Batch** | **HA(ug/mL)** | **Type** | **origin** | **Viral Strain** |
| --- | --- | --- | --- | --- | --- |
| Arepanrix | AFLPA328AA | 15 | Vaccine | Canada | X179A |
| Arepanrix | AFLPA359AA | 15 | Vaccine | Canada | X179A |
| Arepanrix | AFLPA373BA | 15 | Vaccine | Canada | X179A |
| Arepanrix | AFLPA319BB | 15 | Vaccine | Canada | X179A |
| Arepanrix | SF1B0454CL | 457 | Bulk | GSK | X179A |
| Pandemrix | AFLSA208A | 15 | Vaccine | GSK | X179A |
| Pandemrix | AFLSA174AA | 15 | Vaccine | France | X179A |
| Pandemrix | AFLSFDA280 | 139 | Bulk | GSK | X179A |
| Pandemrix | AFLSA167AB | 15 | Vaccine | Sweden | X179A |
| Pandemrix | AFLSA097AA | 15 | Vaccine | Sweden | X179A |
| Pandemrix | AFLSA096AA | 15 | Vaccine | Sweden | X179A |

### Mass spectrometry (MS)

MS was performed on Trypsin/Lys-C (Promega), and Chymotrypsin (Promega) digests of Pandemrix and Arepanrix samples (6.5 μg each) and detailed elsewhere 30. Raw mass spectra of each vaccine digest (trypsin and chymotrypsin separately) were analyzed using a combination of Preview and Byonic v.1.4 spectral analysis software (Protein Metrics, CA, USA). Complementary approaches were pursued. Spectrograms derived from each vaccine digest were mapped to (i) a concatenated FASTA file containing the canonical proteomes for five influenza viral strains: A/California/07/2009 (H1N1), NYMC X-181A (identical to NYMC X-181), NYMC X-179, NYMC X-179A, and A/Puerto Rico/8/1934, and (ii) an in-silico mutant peptide library artificially generated from X-179A that included all possible single amino acid substitutions in the five most frequent flu proteins out of 14,894 unique proteins. Additional validation was done using typical False Discovery Rate (FDR) by including common contaminants and sequence reverses in the FASTA database for all searches. Data was qualified down to a 1% FDR level for proteins. Further fragments that passed a threshold of log probability of 2 and were present four or more times in at least one vaccine digest were considered for further analysis.

### Statistical Analysis

After the MS vaccine digest files were aligned to the reference library, the mass spectra were exported as CSV files from Byonic v.1.4 spectral analysis software (Protein Metrics, CA, USA) and then processed through custom R and Python scripts. Using the Biopython library, the protein sequences for the five reference flu proteins used in the MS processing were retrieved. For each reference flu protein, all corresponding peptides were extracted from the MS database and compared to the reference protein sequence at each motif. From this comparison, the frequency of mutated and wild-type amino acids relative to the X-179A derived viral strain was characterized at each position in each protein in each vaccine batch. Mutation proportions were then computed for each position. After trypsin and chymotrypsin data from the same vaccine batch were merged, a two-tailed Student’s t-test was used to determine significant mutation proportions between Pandemrix and Arepanrix batches, thus pinpointing the specific mutations that significantly distinguished Arepanrix and Pandemrix vaccines.

### DQ0602 binding

DQ0602 is strongly associated with narcolepsy susceptibility, with further increased susceptibility being observed if an individual is homozygous for DQ0602 ^37^. Further, as mentioned above, in addition to a strong DQ0602 association, the T-cell receptor alpha loci are also strongly associated with narcolepsy^10^. Using a combination of in-vitro and in-silico methods, we sought to assess if mutated motifs (see table 3) could modify the DQ0602 peptide-binding register and subsequently, the overall binding affinity to DQ0602. The *in-vitro* binding has been described in detail in Luo et al^32^.

#### In silico binding

The DQ0602 peptide binding prediction algorithm (http://tools.iedb.org/mhcii/) was used to assess if mutated motifs derived from the mass spectroscopic readouts changed DQ0602 peptide-binding registers. Consequently, three 20mer peptide variants derived from the X179 strain, wild type pH1N1, and mutated motif from the vaccines were used to query the DQ0602 binding prediction server^38^. The 20 mer peptide stretches surrounding the mutated motif (10 aa upstream and 10 aa downstream of the mutated motif) or the homologous position in the reference strains were constructed using custom python scripts.

### Tetramer analysis

For a few selected peptides where mutations were found to alter T cell reactivity when presented by DQ0602 potentially, and when these could be hypothesized to explain why Pandemrix could have been more narcolepsy triggering than Arepanrix, DQ0602 peptide tetramers were created and tested in a few DQ0602 narcolepsy and controls subjects as described in Luo et al^32^.

## Results

### Characterization of Protein Content in Arepanrix and Pandemrix

Each vaccine lot was digested with trypsin and chymotrypsin and then subjected to MS characterization, thereby ensuring high coverage of the representative protein content. Mean coverage for the influenza proteins was 80.5% in Pandemrix versus 71.1% in Arepanrix (supplementary figure 1). We observed highly similar mean global proportions of influenza (Pandemrix 60.2 %; Arepanrix 59.4 %), chicken (Pandemrix 31.7; Arepanrix 31.4%) and bovine proteins (Pandemrix 7.9%; Arepanrix 9.1%) in these vaccines (figure 1A). Such results are consistent, given that both vaccines are produced from the same parent NYMC X-179A reassortant virus consisting of PR8 backbone and pH1N1 surface proteins. Bovine proteins are likely reflecting deoxycholate solubilization, as this compound is isolated from bovine gallbladder extracts.

**Figure 1:**
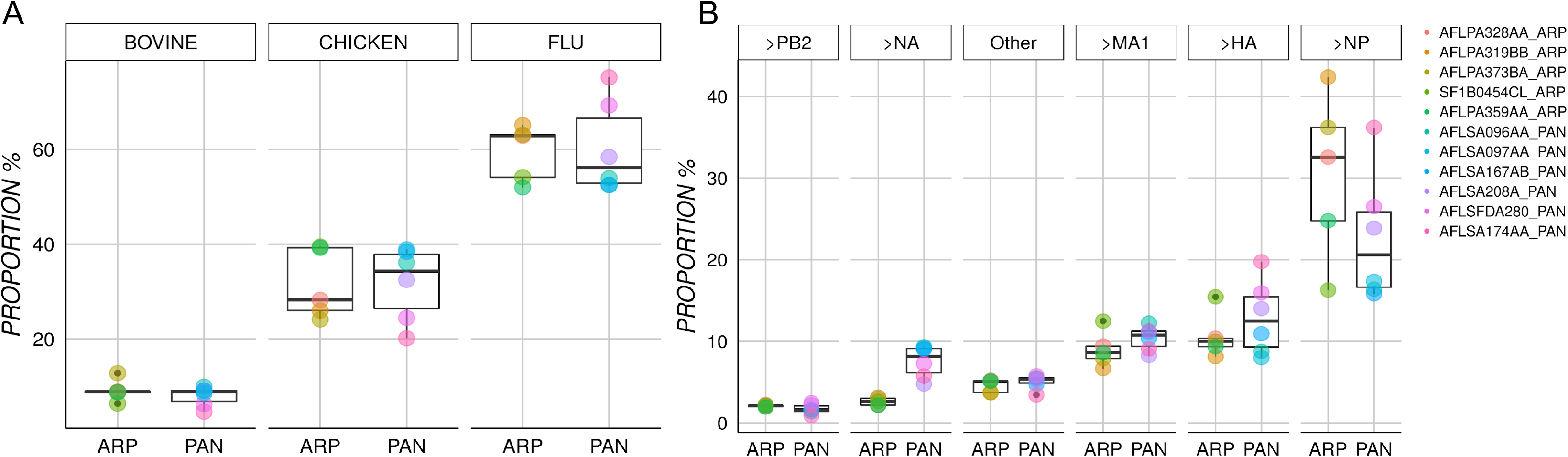
Protein composition in Pandemrix versus Arepanrix. 1A-Boxplots for each type of protein, the number of corresponding peptide fragments was cumulatively counted across all the batches for each vaccine and globally classified as Influenza (FLU), chicken, and bovine origin. 1B-This figure shows the relative proportion or concentration of each protein type within each vaccine for peptide fragments classified as influenza proteins.

In Pandemrix, influenza neuraminidase (NA) was significantly overrepresented (7.56 % vs 2.64%; p=0.0012), hemagglutinin (HA) was moderately increased (12.9% vs 10.6%, p = 0.3), while nucleoprotein (NP) was underrepresented (22.69% vs 30.44% p=0.2) (Figure 1B). The proportion of other influenza proteins like matrix protein (MA1), polymerase PB2, polymerase PA, polymerase PB1 were similar in both Pandemrix and Arepanrix (Table 2). Although the global proportion of non-viral proteins were similar in both Arepanrix and Pandemrix, we observed that chicken proteins such as ApoB, Vitellogenin-2, Glucose-6-phosphate isomerase, Ovalbumin, Ezrin, and Annexin A2 were significantly overrepresented (p<0.05) in Arepanrix (Table 2). The bovine proteins such as tubulin alpha-1B were significantly overrepresented in Pandemrix, while Junction plakoglobin was significantly overrepresented in Arepanrix.

**Table 2:** The protein content of arepanrix and pandemrix

| Protein | Organism | Pandemrix | Arepanrix | CI95 U±L | Statistic | p-value |
| --- | --- | --- | --- | --- | --- | --- |
| Nucleoprotein NP | PR8 | 22.69% (1600) | 30.44% (1092) | 20.8 ± -5.3 | 1.389 | 0.2044 |
| Hemagglutinin HA | pH1N1 | 12.9% (894) | 10.66% (420) | 2.9 ± -7.3 | -1.003 | 0.3436 |
| Matrix Protein 1 MA1 | PR8 | 10.39% (724) | 9.01% (357) | 1.4 ± -4.1 | -1.197 | 0.2709 |
| Neuraminidase NA | pH1N1 | 7.56% (533) | 2.64% (99) | -2.9 ± -7 | -5.997 | **0.0012** |
| polymerase PB2 | PR8 | 1.7% (109) | 2.09% (80) | 1 ± -0.2 | 1.655 | 0.1528 |
| polymerase PA | pH1N1 | 1.81% (124) | 1.34% (56) | 0.5 ± -1.5 | -1.120 | 0.3003 |
| polymerase PB1 | pH1N1 | 1.4% (91) | 1.33% (52) | 0.6 ± -0.7 | -0.266 | 0.7985 |
| nonstructural protein 1 | PR8 | 0.52% (34) | 0.45% (18) | 0.1 ± -0.3 | -0.821 | 0.4404 |
| nuclear export protein | PR8 | 0.11% (10) | 0.26% (11) | 0.4 ± -0.1 | 1.605 | 0.1496 |
| Apolipoprotein B | *Gallus gallus* | 0.16% (11) | 2.07% (87) | 3.2 ± 0.6 | 3.965 | **0.0160** |
| Vitellogenin-2 | *Gallus gallus* | 0.12% (8) | 1.15% (47) | 1.5 ± 0.6 | 5.772 | **0.0024** |
| Glucose-6-phosphate isomerase | *Gallus gallus* | 0.49% (35) | 0.87% (31) | 0.7 ± 0 | 2.684 | **0.0442** |
| Ovalbumin | *Gallus gallus* | 0.39% (28) | 0.85% (33) | 0.8 ± 0.1 | 2.872 | **0.0200** |
| Annexin A2 | *Gallus gallus* | 0.55% (37) | 0.73% (28) | 0.4 ± 0 | 2.382 | **0.0434** |
| Ezrin | *Gallus gallus* | 0.58% (40) | 0.73% (28) | 0.3 ± 0 | 2.754 | **0.0246** |
| Junction plakoglobin | *Bos taurus* | 0.15% (9) | 0.69% (23) | 1.1 ± 0 | 2.586 | **0.0496** |
| Tubulin alpha-1B chain | *Bos taurus* | 0.88% (60) | 0.54% (21) | -0.1 ± -0.6 | -3.212 | **0.0147** |

### Differential mutation proportions among Arepanrix and Pandemrix

A comparison of mutations in Pandemrix and Arepanrix uncovered 17 significantly interesting site-specific differences in relation to reference X-179A strain. 4 HA motifs were represented differently among the vaccines. Most significantly, the HA 146 (N > D) residue, which is close to a receptor binding site that interacts with human respiratory epithelial cells to initiate infection, was deamidated nearly six times more in Arepanrix (59.7%) than Pandemrix (10.7%). This mutation was by far the most significant difference among the two vaccines (p =4.4×10^−6^), confirming previous observations that the difference in HA 146 was an apparent outlying mutation between Arepanrix and Pandemrix^30^. The other mutated residues in hemagglutinin included HA 314 (P > Q, p=5.7×10^−4^), HA 482 (F >Y, p=8.4×10^−3^), and HA 420 (R > I, p=4.2×10^−2^), all significantly enriched at least 3-fold in Arepanrix (Table 3, Figure 2A).

**Table 3:** Significantly different mutated motifs as compared to X179A strain in six batches of Pandemrix and five batches of Arepanrix

| Protein Pos<br>(initial > mut) | P-value | T-stat (CI U ± L) | Mean Proportion % |  | DQ Binder |
| --- | --- | --- | --- | --- | --- |
|  |  |  | Arepanrix | Pandemrix |  |
| HA 146 (N > D) | 4.40E-06 | 10.2 (60 ± 38) | 59.7 | 10.7 | Yes‡ |
| NP 11 (E > Q) | 2.80E-05 | 7.9 (11.1 ± 6.2) | 10.5 | 1.9 | No |
| NP 257 (T > A) | 5.20E-04 | -6.8 (-4 ± -8.5) | 0.3 | 6.5 | Yes† |
| HA 314 (P > Q) | 5.70E-04 | 7.3 (55.6 ± 27) | 44.2 | 2.9 | No |
| MA 192 (N > D) | 2.61E-03 | 4.2 (7.6 ± 2.2) | 10.2 | 5.3 | No |
| NP 424 (T > I) | 3.65E-03 | -5.1 (-3.6 ± -10.8) | 0.1 | 7.3 | Yes* |
| M1 49 (R > I) | 3.90E-03 | 4.6 (76.8 ± 23.4) | 79.1 | 29 | No |
| HA 482 (F > Y) | 8.24E-03 | 4.3 (51.2 ± 12.6) | 38.5 | 6.6 | No |
| NP 469 (E > D) | 1.13E-02 | 3.9 (12.9 ± 2.6) | 11.5 | 3.8 | No |
| NP 432 (N > D) | 1.28E-02 | 3.1 (13.6 ± 2.1) | 17.2 | 9.3 | No |
| M1 91 (N > D) | 1.66E-02 | 3.1 (13.2 ± 1.8) | 17.7 | 10.3 | No |
| M1 85 (N > D) | 1.78E-02 | -3.4 (-2.8 ± -19) | 23.5 | 34.4 | No |
| M1 99 (L > M) | 2.12E-02 | 2.8 (11 ± 1.1) | 10.4 | 4.3 | No |
| NP 321 (N > D) | 2.14E-02 | 3.3 (41.8 ± 5.3) | 29.2 | 5.7 | No |
| M1 84 (L > P) | 2.24E-02 | -2.9 (-1.2 ± -11.6) | 14.9 | 21.3 | No |
| NP 423 (T > R) | 3.19E-02 | -2.9 (-0.5 ± -7.4) | 0 | 4 | Yes* |
| HA 420 (R > I) | 4.42E-02 | 2.8 (45 ± 1) | 24.2 | 1.3 | No |

\*Reduced binding in Pandemrix, but specific of pandemrix. †Same binding, different TCR facing residues. ‡Unlikely to be involved, but present in pH1N1.

**Figure 2:**
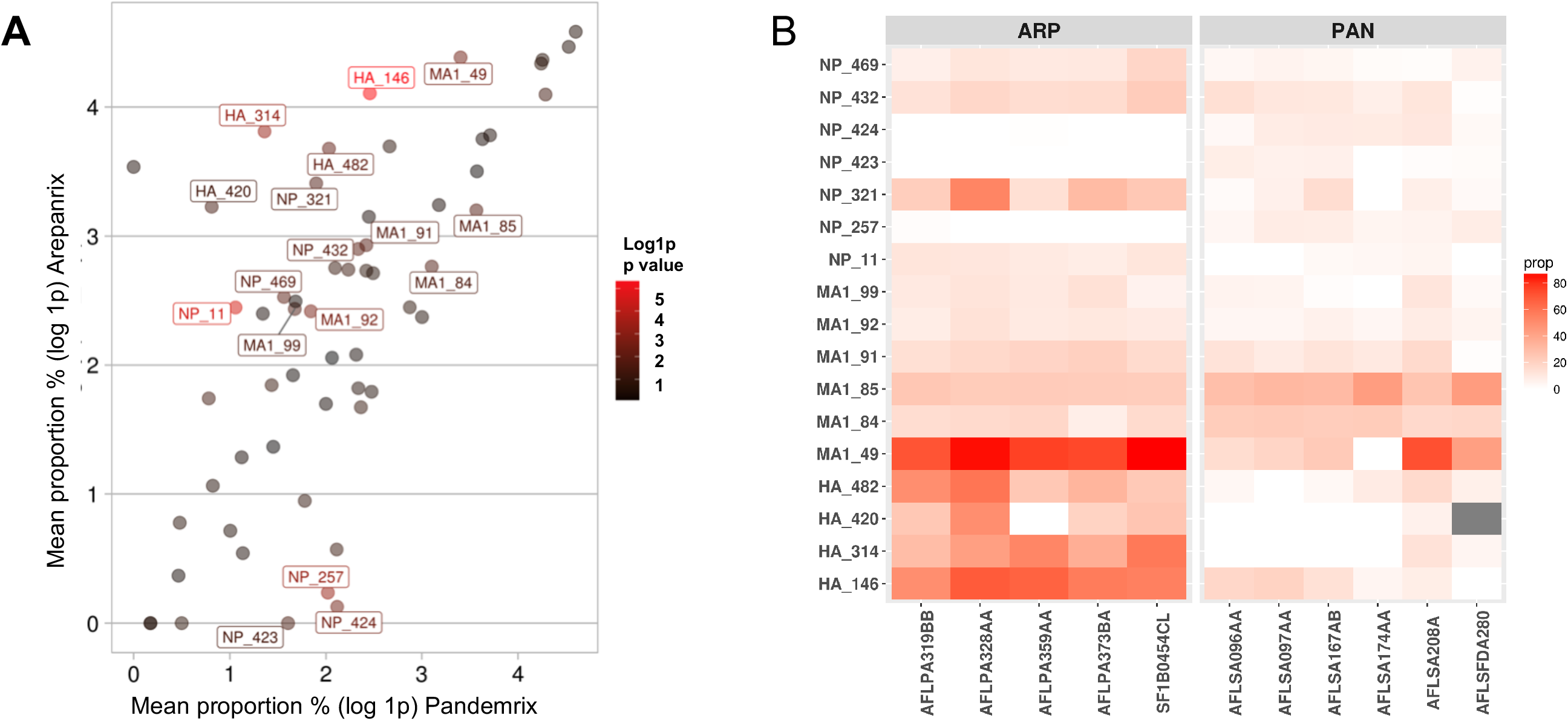
Mean mutation proportion in Arepanrix versus Pandemrix. (2A) Each data point represents a specific mutated amino acid. This scatterplot maps the log-transformed mean mutation proportion in Pandemrix on the x-axis against log-transformed mean mutation proportion in Arepanrix on the y-axis. Calculated from a 2-tailed Student’s t-test, the p-value of each data point indicates how significantly different the mutation is between vaccines. As data points are colored based on their p-value, the redder on the gradient scale, the more significant the mutation is in regards to differentiating Pandemrix and Arepanrix. (2B) Heatmap is indicating the actual mutational proportion with the y-axis showing the mutated positions and x-axis the vaccine lots.

NP, which is derived from PR8 and not pH1N1 sequence, also accumulated mutations that differentiated the two vaccines. For instance, the NP 11 (E > Q, p=2.8e-5) residue mutated nearly 5 times more in Arepanrix (10.5%) than Pandemrix (1.9%), while deamidations were frequently enriched in Arepanrix at NP 321 (N > D, p=0.02) and NP 432 (N > D, p=0.01). It should be noted that some NP mutations occurred more frequently in Pandemrix than Arepanrix; the most significant of these include the following residues: NP 257 (T > A, p=5.2×10^−4^), NP 423(T > R, p=3.1×10^−2^) and NP 424 (T > I, p=3.6×10^−3^) (Table 3 and full list of mutations per batch in Supplementary Table 6).

Matrix protein 1, also derived from PR8, was observed to have a differential accumulation of mutations between Arepanrix and Pandemrix. The most significant differences include the residue M1 49 (R > I, p=3.9e-3) as well as adjacent motifs deamidated at M1 91 (N >D, p=1.6 ×10^−2^), M1 92 (N >D, p=2.6×10^−3^) and M1 99 (L > M, p=2.1e-2), that were all over represented in Arepanrix. Meanwhile, two adjacent motifs at M1 84 (L > P, p=2.2 ×10^−2^) and M1 85 deamidation (N > D, p=1.7 ×10^−2^) were enriched in Pandemrix (Table 3).

Statistical comparison of mutation proportions revealed a general trend in increased differential mutations across in Arepanrix compared to Pandemrix and, as shown in Figure 2. Of interest, statistically significant (p-value < 0.05) mutations, such as HA 146, HA 314, and NP 11, typically occurred more often in Arepanrix than Pandemrix.

### DQ0602 binding of mutated motifs in Arepanrix and Pandemrix

We next sought to determine whether these mutated and enriched motifs (Table 3) in either Pandemrix or Arepanrix modified their HLA binding registers and subsequently their overall propensity to bind narcolepsy associated DQ0602 allele. A combination of in-vitro and in-silico methods was adopted to address this question. We synthesized 15 mer peptide stretches overlapping by 4mers using the most abundant viral proteins (HA, NA, NP, M1 & PB2) from reference strains (X179A and wild type pH1N1) as a template. These 15 mer peptides were used to quantify DQ0602 binding affinities relative to a known EBV derived 15 mer peptide that is a strong binder^32^. In this way, a database of experimentally derived peptide binders from the abundant viral proteins was built. Next, we compared the mutated motifs (Table 3) from the vaccines to the reference motifs already tested for DQ0602 binding.

Among the mutations described in table 3, six of the variant motifs changed the DQ0602 binding register. First, we confirmed that the previously described HA 146 (N to D) mutation enriched in Arepanrix changed peptide-binding register to bind strongly to DQ0602 (N allele 4.4 %, D allele 53.47 %). Second, we identified five novel mutations that changed DQ0602 binding registers. NP 257 (T > A) was enriched in Pandemrix and changed the binding register to DL[A]FLARSA (A allele 4.8%) from the reference DL[T]FLARSA (T allele 15.7%), this change is again projected to increase the binding affinity to DQ0602. Other mutations that modified DQ0602 binding registers are NP 423, HA 420, MA1 49 (see Supplementary Table 1 & 2). Considering their differential abundance in Pandemrix versus Arepanrix and expected changes in binding register, we determined that only four mutations i.e., HA 146, NP 423, NP 424, and NP 257, have the potential to explain differential effects of these vaccines on narcolepsy.

### Tetramer studies of 4 mutated motifs that could have impacted narcolepsy risk

Our analysis projected that four Pandemrix enriched mutations (i.e., HA 146, NP 423, NP 424, and NP 257, see supplementary table 1 & 2) within the DQ0602 binding registers could influence narcolepsy susceptibility. We thus conducted DQ0602 tetramer screens of the mutated motifs in expanded PBMCs as described previously^32^ split into three conditions (i.e. stimulated with Arepanrix, Pandemrix, or specific mutated peptide) to identify cognate mutated peptide-specific CD4+ T cells. Six narcolepsy cases (5 post Pandemrix, one recent onset) and 4 DQ0602 control cases (all vaccinated with Pandemrix in 2009-2010) described in Luo et al. .^32^ were selected for this screen. While there appeared to be sporadic reactivity to some mutated motifs, we did not observe any significant differences in the frequencies of tetramer specific CD4 T cells in narcolepsy cases vs. controls (supplementary table 4 and supplementary table 5). In comparison to immunodominant motifs we identified in our prior screens in these same patients, these epitopes^18^ were considered insignificant.

## Discussion

This study extends the Jacob et al. report^30^ where only single batches of Arepanrix and Pandemrix were analyzed and presents a detailed characterization of the mutational burden and protein content of 5 Arepanrix and 6 Pandemrix batches. Mean coverage of the mass spectrometric characterization of influenza proteins, while still at high 71.1 % in Arepanrix and 80.5 % in Pandemrix, was slightly less than what Jacob et al. .^30^. The sampled lots were actual vaccine doses used during the 2009 pandemic influenza vaccination campaign in Northern Europe and Canada, except for two lots that were monovalent bulks and sourced directly from GSK (see Table 1). Not surprisingly, considering that both Arepanrix and Pandemrix were derived from NYMC X-179A^26,30^, the mean global proportions of Influenza, chicken and bovine proteins were comparable between the two vaccines (figure 1). This finding agrees with Jacob et al. ^26^, where similar global proportions were observed. The four main influenza proteins in order of abundance were NP, HA, M1, NA, and PB2 (table 2), which is consistent with other studies^26^, while PB1, NS1 and nuclear export protein (NEP) were only present at low concentrations (< 1%) in both the vaccines.

We found that NA was enriched three-fold in Pandemrix as compared to Arepanrix, while NP was underrepresented in Pandemrix as compared to Arepanrix. This finding conflicts with reports of Jacob et al. .^26^, Vaarala et al^33^. and Ahmed et al^34^., all of which observed an overrepresentation of NP content in Arepanrix. The main limitation in these studies mentioned earlier is however that only one representative batch of each vaccine was characterized by mass spectrometry; in this study, we characterized 6 Pandemrix and 5 Arepanrix batches, using both trypsin and chymotrypsin protein digests to increase our protein coverage thus the finding may be more reliable. However, we did not perform any enrichment before MS characterization, which may have influenced our current results.

As recently reported, we found that vaccine strains, like wild type virion infecting hosts, mutate in culture, and this leads to divergences in vaccine viral sequences in different vaccines or across vaccine batches. Differences may thus depend on how often the manufacturer reuses the primary NYMC strain versus continuing to amplify isolates from their own egg cultures for future propagation. As an example, Skowronski et al. found that H3N2 reassortant vaccine strains had mutated in key antigenic residues, likely contributing to reduced efficacy in 2012-2013^39^. Similarly, Jacob et al., conducting Mass Spectrometry (MS) characterization of X-179A derived pH1N1 vaccines, 2009 Pandemrix and 2010 Arepanrix, discovering that a specific HA mutation, N146D, had accumulated in Arepanrix, distinguishing the two antigens^30^ (limitation in the Jacob et al. study was that only a representative batch of actual Pandemrix vaccine (batch DFLSA014A) and bulk Arepanrix (batch SF1B0454Cl) was studied and compared. Further, the Arepanrix lot was a lot that had never been used and had been prepared one year after the pandemic (2010). In this study, we could confirm the dominance of N146D in Arepanrix but not Pandemrix across all lots. Interestingly, this mutation conferred higher growth and was selected in subsequent pH1N1 strains X-181^26^. It may thus be that the mutation accumulated in Arepanrix but not Pandemrix because of differences in Arepanrix culture procedures.

As it was conceivable that the N146 sequence found in Pandemrix and wild type H1N1 but not Arepanrix was essential to explain narcolepsy susceptibility, we further examined binding of both N146 and D146 peptides on DQ0602 molecules, confirming in vitro prediction indicating that 146 binds with lower affinity to DQ0602, another factor that could contribute to different susceptibility. Using DQ0602 tetramers for sequences; however, we found that very few T cells recognized these peptide sequences in both narcolepsy and control subjects, making it unlikely to be of significance in narcolepsy pathophysiology. Similar to the study of HA N146D, we also studied tetramers for NP T424I, NP T423I, and NP T257A, three other mutations that are much more abundant in Pandemrix versus Arepanrix (table 1) and were predicted to bind DQ0602 with an equivalent affinity (supplementary table 1). In vitro binding studies indeed found that all these peptides bound DQ0602 with high affinity. However, tetramer studies in narcolepsy and controls did not support abundance for T cells recognizing these epitopes making it unlikely to be functionally important.

With recent results suggesting that a potential mimic of HCRT^NH2^ is pHA^273-287^, we also carefully examined frequency and sequence variation within this segment, present in both wild type pH1N1, X-179A and X181, but could not find any mutation or difference in frequency across vaccines, making it unlikely composition difference at this level explain differential narcolepsy risk. It is nonetheless interesting to note that pHA^273-287^ contains N at position 273 (predicted to bind DQ0602 in P1) and that this residue is partially glycosylated^30^, a modification that could make a difference in B cell reactivity and perhaps epitope processing and presentation to T cells. A future direction would be to profile in detail the post-translational modifications in these motifs across the two vaccines. For instance, possible deamidation in the MHC binding pocket could alter the DQ0602 binding register and glycosylation in or flanking the pHA^273-287^ or in other cases, PTMs such as N-linked or O-linked glycosylation have also been shown to protect epitope cleavage sites, prevent efficient antigen processing, or influence recognition by cognate T-cell receptors^40^. Glycosylation patterns in key residues across vaccines are, therefore, also of interest in the context of DQ0602 binding and narcolepsy susceptibility.

In summary, characterization of the mutation load of several Pandemrix and Arepanrix lots revealed extensive differences in influenza mutation frequencies when compared across the vaccines and also in relation to the parent vaccine strain X179A. We also did not find any single mutations within the most likely culprit mimic sequence that may trigger autoimmunity, pHA^273-287^ in this study. Future research exploring double mutations or PTMS in this region (pHA^273-287^) and additional flanking regions are underway in our laboratory and could yield additional answers. Nonetheless, as identified in this study, the vaccine composition is complex and diverse, and thus unrecognized differences could still play a role. As an example, it is notable that a significant portion of spectrograms in these vaccines does not map to any known protein sequence, thus our search for differences that could be involved in narcolepsy triggering was in no way exhaustive. Beside vaccine differences, it may also well be that difference in past or concomitant microbial exposure of the Canadian (vaccinated with Arepanrix), and northern European population (vaccinated with Pandemrix) could be primarily involved, for example, timing of concomitant pH1N1 infection, past H3N2 infections, or presence of specific streptococcal A infections, as suggested by other studies^41^.

## Conflict of Interest

EM currently receives funding from Jazz Pharmaceutical, and EM previously received funding from GlaxoSmithKline for the study of the immunological basis of post-Pandemrix-narcolepsy. Conduct for these studies was supervised by and reported to the European Medical Agency. GlaxoSmithKline holds the patent for Pandemrix. Funding from these two sources did not support the research published in the manuscript. Besides, a provisional patent on a potential DQ0602 hemagglutinin flu epitope sequence cross-reactive with hypocretin was filed by GSK and Stanford with EM as one of the inventors, but the patent was subsequently abandoned when the publication of De la Herran-Arita et al^42,43^ was retracted. All other authors report no conflict of interest relevant to this manuscript.

---

**Supplementary Figure 1:**
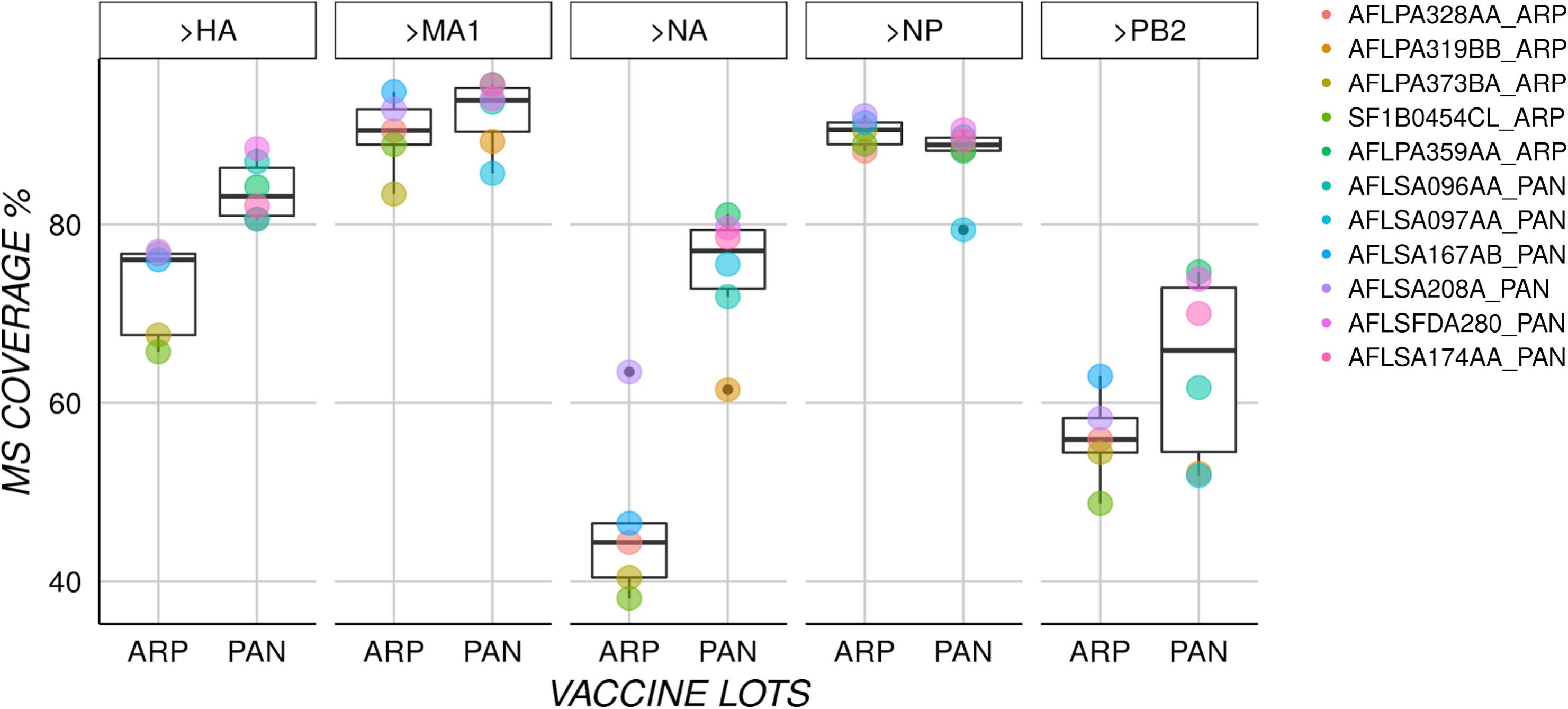
The percentage coverage of the main influenza viral proteins as characterized by mass spectrometry across different vaccine lots derived from Pandemrix and Arepanrix post digestion with trypsin and chymotrypsin.

## Notes

### Competing Interest Statement

The authors have declared no competing interest.

