## Supplementary figure 2 for "Mass Spectrometric Characterization of Narcolepsy-Associated Pandemic 2009 Influenza Vaccines"

### Supplementary Figure 2 : Raw FACS staining plots of mutant peptide tetramers in

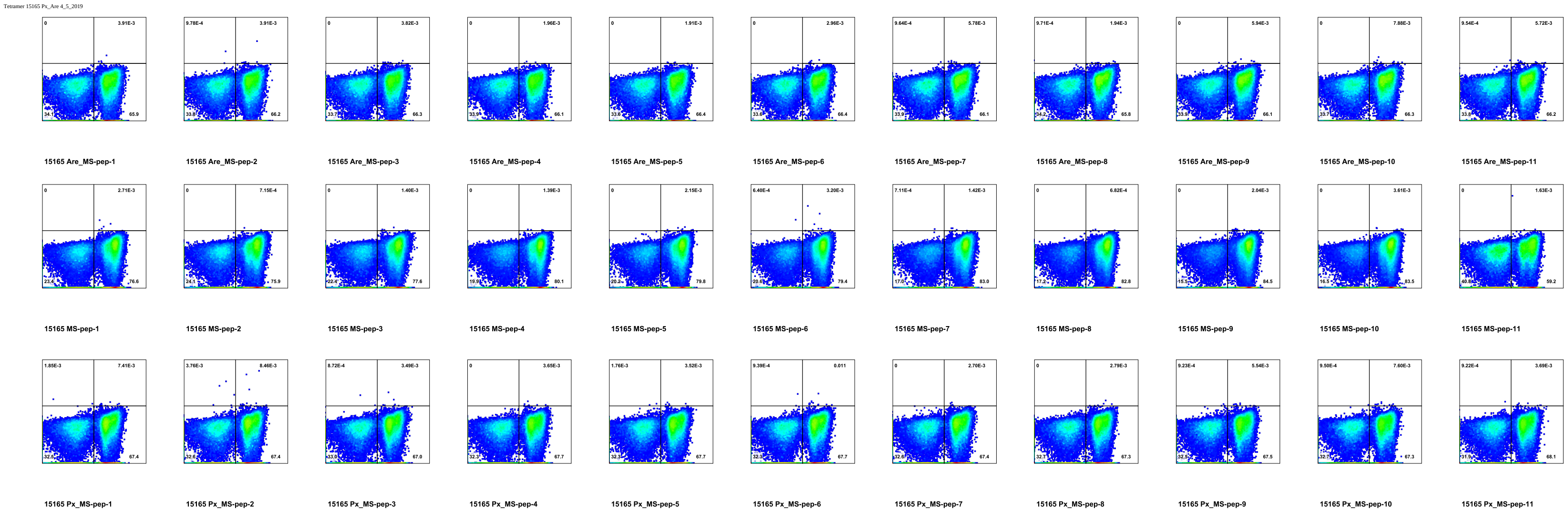

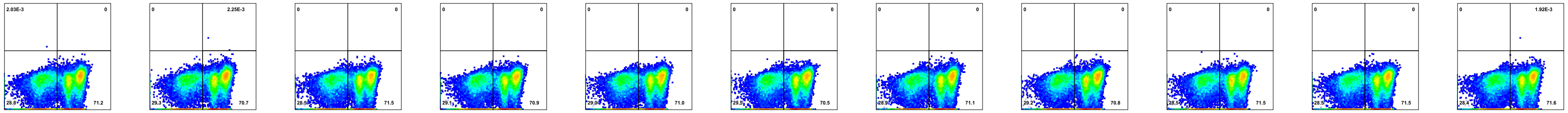

16021_9d Are_MS-pep-1 | 16021_9d Are_MS-pep-2 | 16021_9d Are_MS-pep-3 | 16021_9d Are_MS-pep-4 | 16021_9d Are_MS-pep-5 | 16021_9d Are_MS-pep-6 | 16021_9d Are_MS-pep-7 | 16021_9d Are_MS-pep-8 | 16021_9d Are_MS-pep-9 | 16021_9d Are_MS-pep-10 | 16021_9d Are_MS-pep-11

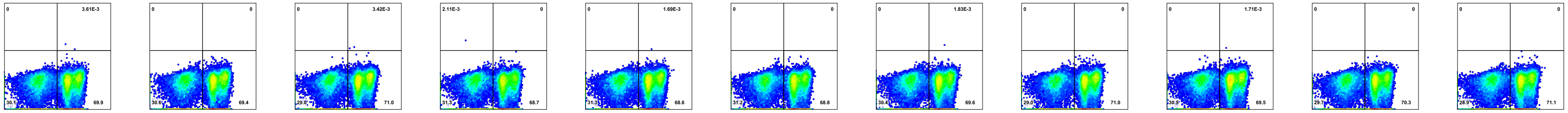

16021_9d MS-pep-1 | 16021_9d MS-pep-2 | 16021_9d MS-pep-3 | 16021_9d MS-pep-4 | 16021_9d MS-pep-5 | 16021_9d MS-pep-6 | 16021_9d MS-pep-7 | 16021_9d MS-pep-8 | 16021_9d MS-pep-9 | 16021_9d MS-pep-10 | 16021_9d MS-pep-11

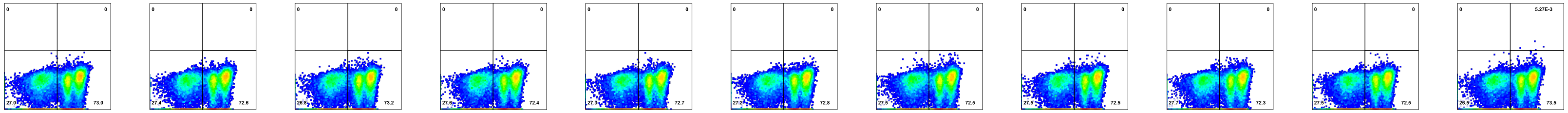

16021_9d Px_MS-pep-1 | 16021_9d Px_MS-pep-2 | 16021_9d Px_MS-pep-3 | 16021_9d Px_MS-pep-4 | 16021_9d Px_MS-pep-5 | 16021_9d Px_MS-pep-6 | 16021_9d Px_MS-pep-7 | 16021_9d Px_MS-pep-8 | 16021_9d Px_MS-pep-9 | 16021_9d Px_MS-pep-10 | 16021_9d Px_MS-pep-11

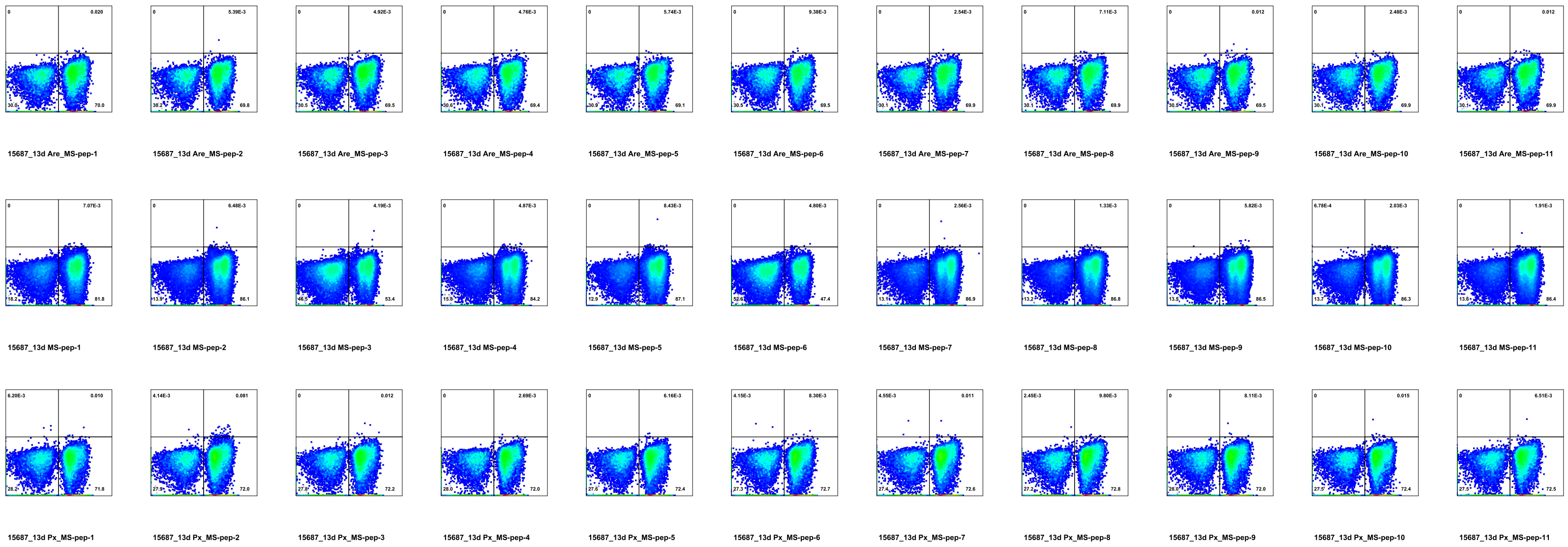
15687_13d Are_MS-pep-1
15687_13d Are_MS-pep-2
15687_13d Are_MS-pep-3
15687_13d Are_MS-pep-4
15687_13d Are_MS-pep-5
15687_13d Are_MS-pep-6
15687_13d Are_MS-pep-7
15687_13d Are_MS-pep-8
15687_13d Are_MS-pep-9
15687_13d Are_MS-pep-10
15687_13d Are_MS-pep-11
15687_13d MS-pep-1
15687_13d MS-pep-2
15687_13d MS-pep-3
15687_13d MS-pep-4
15687_13d MS-pep-5
15687_13d MS-pep-6
15687_13d MS-pep-7
15687_13d MS-pep-8
15687_13d MS-pep-9
15687_13d MS-pep-10
15687_13d MS-pep-11
15687_13d Px_MS-pep-1
15687_13d Px_MS-pep-2
15687_13d Px_MS-pep-3
15687_13d Px_MS-pep-4
15687_13d Px_MS-pep-5
15687_13d Px_MS-pep-6
15687_13d Px_MS-pep-7
15687_13d Px_MS-pep-8
15687_13d Px_MS-pep-9
15687_13d Px_MS-pep-10
15687_13d Px_MS-pep-11

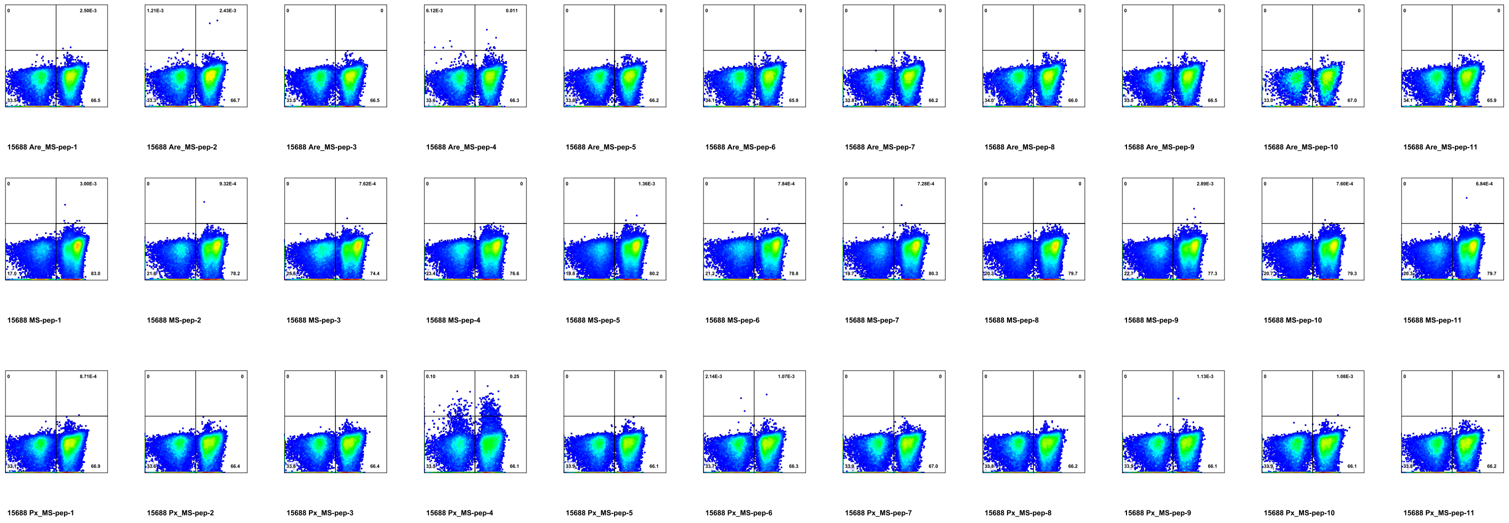
15688 Are_MS-pep-1
15688 Are_MS-pep-2
15688 Are_MS-pep-3
15688 Are_MS-pep-4
15688 Are_MS-pep-5
15688 Are_MS-pep-6
15688 Are_MS-pep-7
15688 Are_MS-pep-8
15688 Are_MS-pep-9
15688 Are_MS-pep-10
15688 Are_MS-pep-11
15688 MS-pep-1
15688 MS-pep-2
15688 MS-pep-3
15688 MS-pep-4
15688 MS-pep-5
15688 MS-pep-6
15688 MS-pep-7
15688 MS-pep-8
15688 MS-pep-9
15688 MS-pep-10
15688 MS-pep-11
15688 Px_MS-pep-1
15688 Px_MS-pep-2
15688 Px_MS-pep-3
15688 Px_MS-pep-4
15688 Px_MS-pep-5
15688 Px_MS-pep-6
15688 Px_MS-pep-7
15688 Px_MS-pep-8
15688 Px_MS-pep-9
15688 Px_MS-pep-10
15688 Px_MS-pep-11

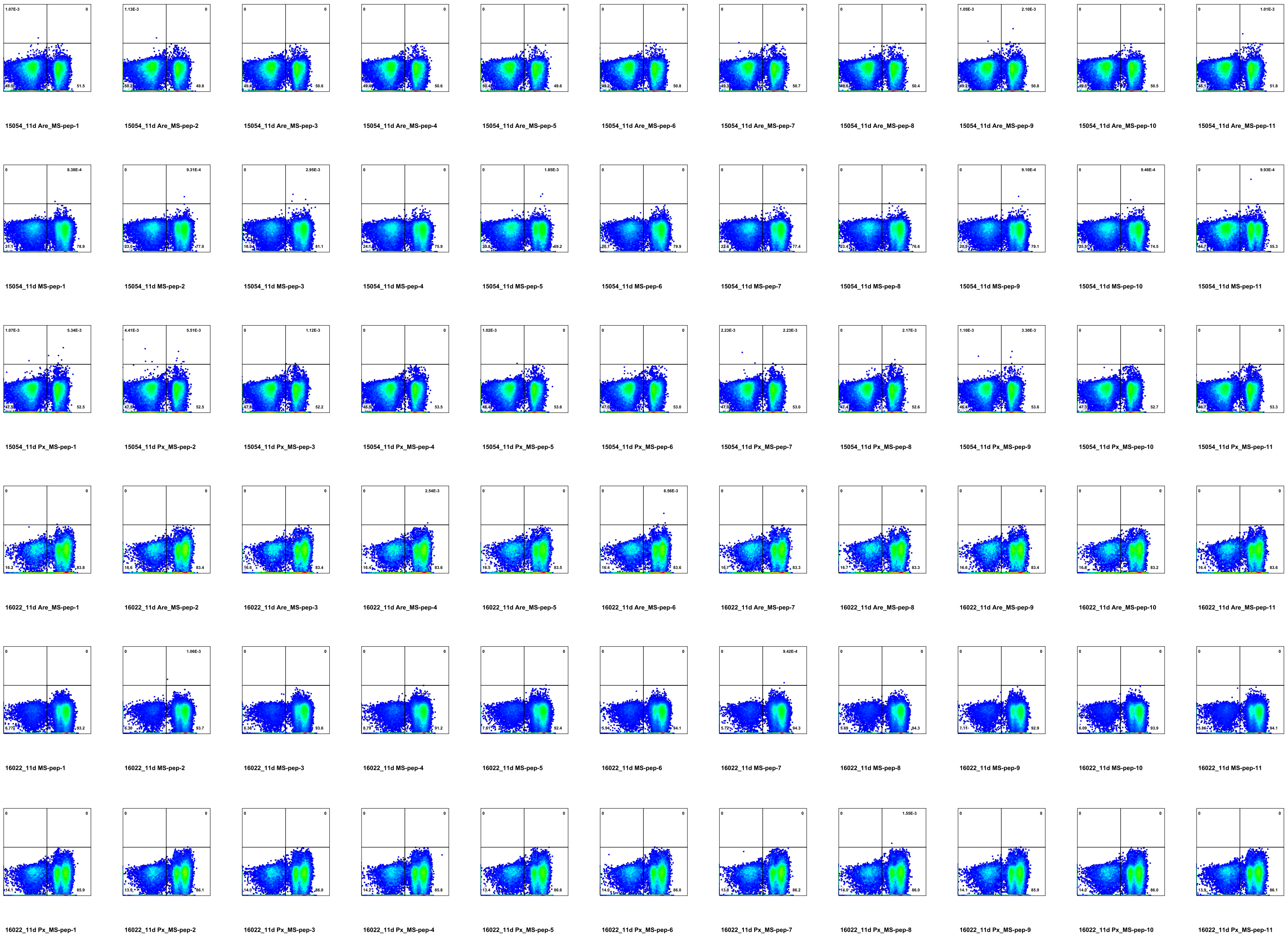
15054_11d Are_MS-pep-1
15054_11d Are_MS-pep-2
15054_11d Are_MS-pep-3
15054_11d Are_MS-pep-4
15054_11d Are_MS-pep-5
15054_11d Are_MS-pep-6
15054_11d Are_MS-pep-7
15054_11d Are_MS-pep-8
15054_11d Are_MS-pep-9
15054_11d Are_MS-pep-10
15054_11d Are_MS-pep-11
15054_11d MS-pep-1
15054_11d MS-pep-2
15054_11d MS-pep-3
15054_11d MS-pep-4
15054_11d MS-pep-5
15054_11d MS-pep-6
15054_11d MS-pep-7
15054_11d MS-pep-8
15054_11d MS-pep-9
15054_11d MS-pep-10
15054_11d MS-pep-11
15054_11d Px_MS-pep-1
15054_11d Px_MS-pep-2
15054_11d Px_MS-pep-3
15054_11d Px_MS-pep-4
15054_11d Px_MS-pep-5
15054_11d Px_MS-pep-6
15054_11d Px_MS-pep-7
15054_11d Px_MS-pep-8
15054_11d Px_MS-pep-9
15054_11d Px_MS-pep-10
15054_11d Px_MS-pep-11
16022_11d Are_MS-pep-1
16022_11d Are_MS-pep-2
16022_11d Are_MS-pep-3
16022_11d Are_MS-pep-4
16022_11d Are_MS-pep-5
16022_11d Are_MS-pep-6
16022_11d Are_MS-pep-7
16022_11d Are_MS-pep-8
16022_11d Are_MS-pep-9
16022_11d Are_MS-pep-10
16022_11d Are_MS-pep-11
16022_11d MS-pep-1
16022_11d MS-pep-2
16022_11d MS-pep-3
16022_11d MS-pep-4
16022_11d MS-pep-5
16022_11d MS-pep-6
16022_11d MS-pep-7
16022_11d MS-pep-8
16022_11d MS-pep-9
16022_11d MS-pep-10
16022_11d MS-pep-11
16022_11d Px_MS-pep-1
16022_11d Px_MS-pep-2
16022_11d Px_MS-pep-3
16022_11d Px_MS-pep-4
16022_11d Px_MS-pep-5
16022_11d Px_MS-pep-6
16022_11d Px_MS-pep-7
16022_11d Px_MS-pep-8
16022_11d Px_MS-pep-9
16022_11d Px_MS-pep-10
16022_11d Px_MS-pep-11

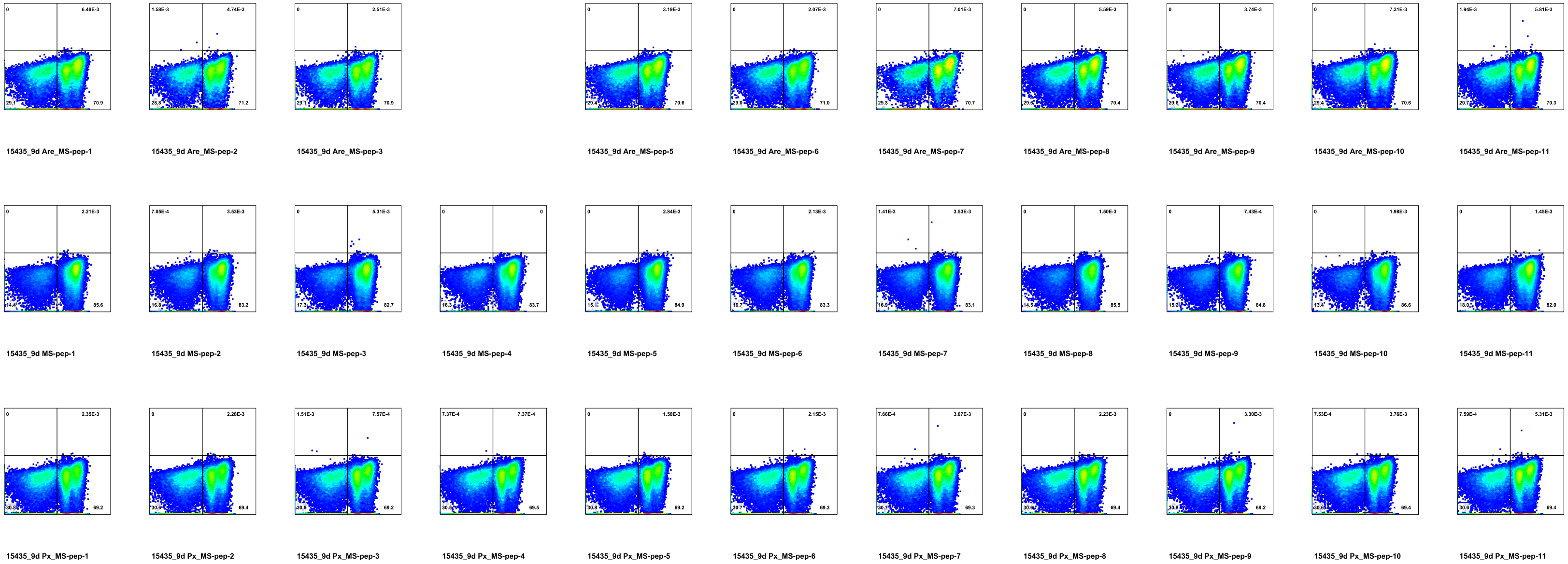
15435_9d Are_MS-pep-1
15435_9d Are_MS-pep-2
15435_9d Are_MS-pep-3
15435_9d Are_MS-pep-5
15435_9d Are_MS-pep-6
15435_9d Are_MS-pep-7
15435_9d Are_MS-pep-8
15435_9d Are_MS-pep-9
15435_9d Are_MS-pep-10
15435_9d Are_MS-pep-11
15435_9d MS-pep-1
15435_9d MS-pep-2
15435_9d MS-pep-3
15435_9d MS-pep-4
15435_9d MS-pep-5
15435_9d MS-pep-6
15435_9d MS-pep-7
15435_9d MS-pep-8
15435_9d MS-pep-9
15435_9d MS-pep-10
15435_9d MS-pep-11
15435_9d Px_MS-pep-1
15435_9d Px_MS-pep-2
15435_9d Px_MS-pep-3
15435_9d Px_MS-pep-4
15435_9d Px_MS-pep-5
15435_9d Px_MS-pep-6
15435_9d Px_MS-pep-7
15435_9d Px_MS-pep-8
15435_9d Px_MS-pep-9
15435_9d Px_MS-pep-10
15435_9d Px_MS-pep-11

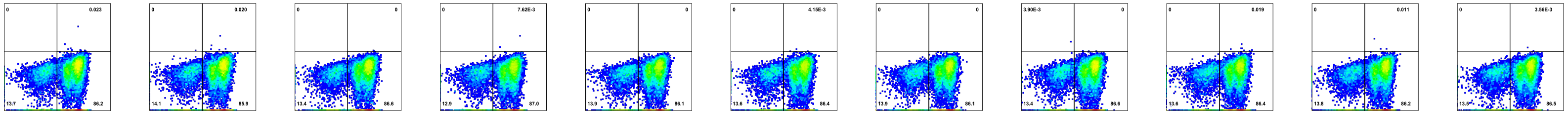

15900 Are_MS-pep-1 | 15900 Are_MS-pep-2 | 15900 Are_MS-pep-3 | 15900 Are_MS-pep-4 | 15900 Are_MS-pep-5 | 15900 Are_MS-pep-6 | 15900 Are_MS-pep-7 | 15900 Are_MS-pep-8 | 15900 Are_MS-pep-9 | 15900 Are_MS-pep-10 | 15900 Are_MS-pep-11

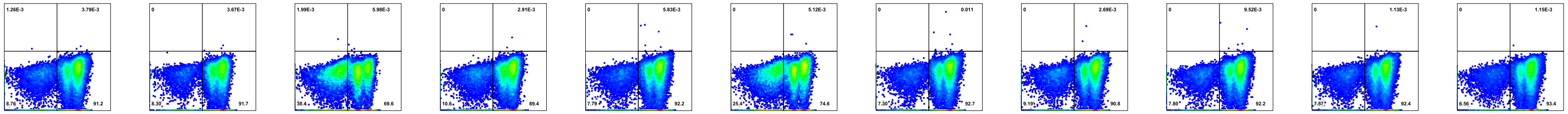

15900 MS-pep-1 | 15900 MS-pep-2 | 15900 MS-pep-3 | 15900 MS-pep-4 | 15900 MS-pep-5 | 15900 MS-pep-6 | 15900 MS-pep-7 | 15900 MS-pep-8 | 15900 MS-pep-9 | 15900 MS-pep-10 | 15900 MS-pep-11

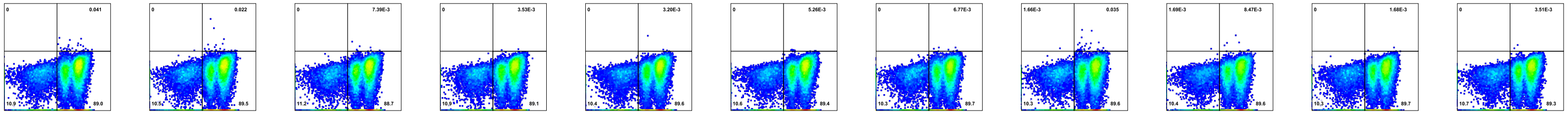

15900 Px_MS-pep-1 | 15900 Px_MS-pep-2 | 15900 Px_MS-pep-3 | 15900 Px_MS-pep-4 | 15900 Px_MS-pep-5 | 15900 Px_MS-pep-6 | 15900 Px_MS-pep-7 | 15900 Px_MS-pep-8 | 15900 Px_MS-pep-9 | 15900 Px_MS-pep-10 | 15900 Px_MS-pep-11

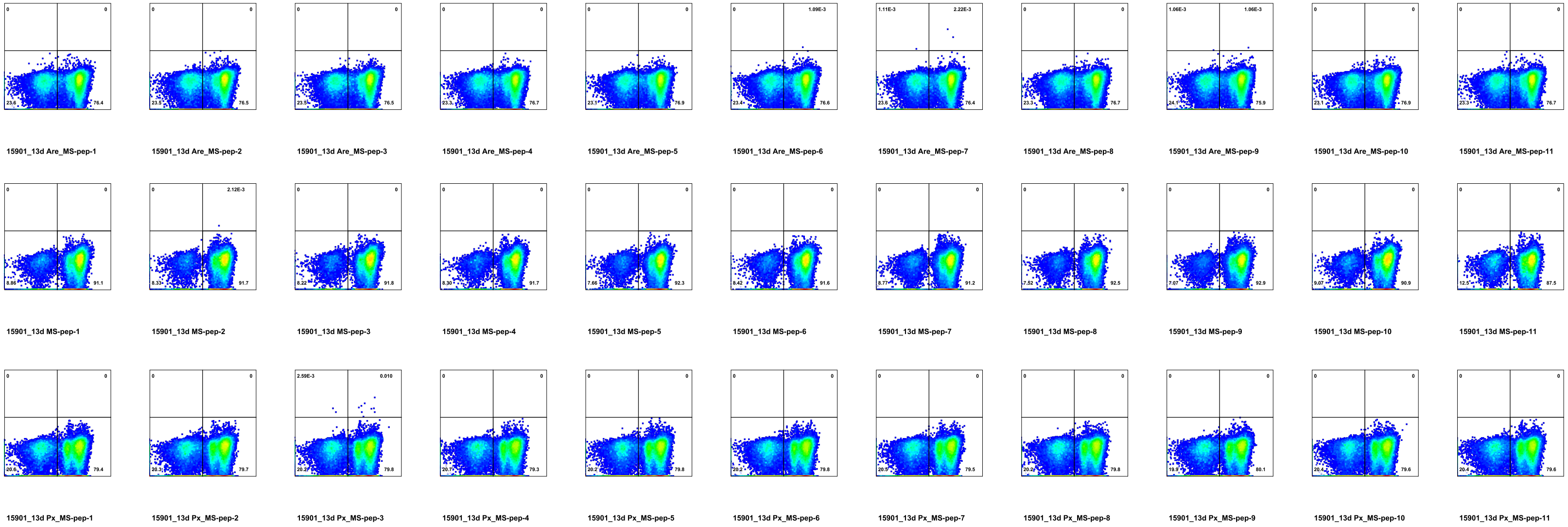
15901_13d Are_MS-pep-1
15901_13d Are_MS-pep-2
15901_13d Are_MS-pep-3
15901_13d Are_MS-pep-4
15901_13d Are_MS-pep-5
15901_13d Are_MS-pep-6
15901_13d Are_MS-pep-7
15901_13d Are_MS-pep-8
15901_13d Are_MS-pep-9
15901_13d Are_MS-pep-10
15901_13d Are_MS-pep-11
15901_13d MS-pep-1
15901_13d MS-pep-2
15901_13d MS-pep-3
15901_13d MS-pep-4
15901_13d MS-pep-5
15901_13d MS-pep-6
15901_13d MS-pep-7
15901_13d MS-pep-8
15901_13d MS-pep-9
15901_13d MS-pep-10
15901_13d MS-pep-11
15901_13d Px_MS-pep-1
15901_13d Px_MS-pep-2
15901_13d Px_MS-pep-3
15901_13d Px_MS-pep-4
15901_13d Px_MS-pep-5
15901_13d Px_MS-pep-6
15901_13d Px_MS-pep-7
15901_13d Px_MS-pep-8
15901_13d Px_MS-pep-9
15901_13d Px_MS-pep-10
15901_13d Px_MS-pep-11

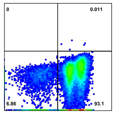
16023_11d Are_MS-pep-1

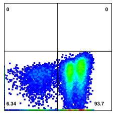
16023_11d Are_MS-pep-2

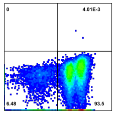
16023_11d Are_MS-pep-3

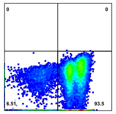
16023_11d Are_MS-pep-4

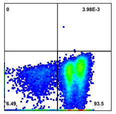
16023_11d Are_MS-pep-5

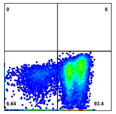
16023_11d Are_MS-pep-6

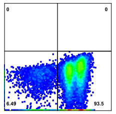
16023_11d Are_MS-pep-7

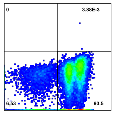
16023_11d Are_MS-pep-8

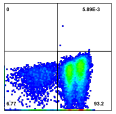
16023_11d Are_MS-pep-9

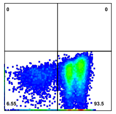
16023_11d Are_MS-pep-10

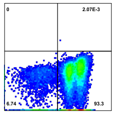
16023_11d Are_MS-pep-11

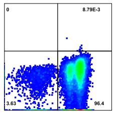
16023_11d MS-pep-1

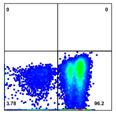
16023_11d MS-pep-2

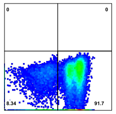
16023_11d MS-pep-3

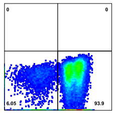
16023_11d MS-pep-4

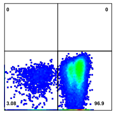
16023_11d MS-pep-5

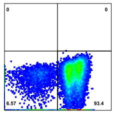
16023_11d MS-pep-6

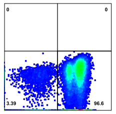
16023_11d MS-pep-7

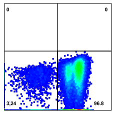
16023_11d MS-pep-8

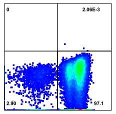
16023_11d MS-pep-9

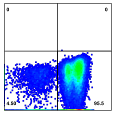
16023_11d MS-pep-10

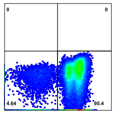
16023_11d MS-pep-11

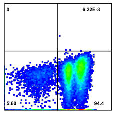
16023_11d Px_MS-pep-1

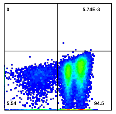
16023_11d Px_MS-pep-2

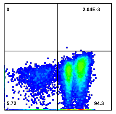
16023_11d Px_MS-pep-3

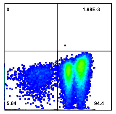
16023_11d Px_MS-pep-4

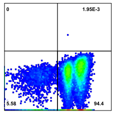
16023_11d Px_MS-pep-5

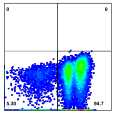
16023_11d Px_MS-pep-6

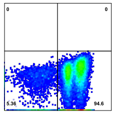
16023_11d Px_MS-pep-7

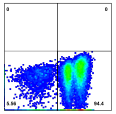
16023_11d Px_MS-pep-8

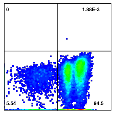
16023_11d Px_MS-pep-9

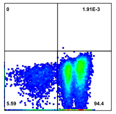
16023_11d Px_MS-pep-10

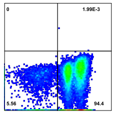
16023_11d Px_MS-pep-11
